# Systems genetics approach uncovers associations between host amylase locus, gut microbiome and metabolic traits in hyperlipidemic mice

**DOI:** 10.1101/2024.02.28.582610

**Authors:** Qijun Zhang, Evan R. Hutchison, Calvin Pan, Matthew F. Warren, Mark P. Keller, Alan D. Attie, Aldons J. Lusis, Federico E. Rey

**Author notes:** Address for Correspondence, Department of Bacteriology, University of Wisconsin-Madison 1550 Linden Dr., Madison, WI 53706.

## Abstract

The molecular basis for how host genetic variation impacts gut microbial community and bacterial metabolic niches remain largely unknown. We leveraged 90 inbred hyperlipidemic mouse strains from the Hybrid Mouse Diversity Panel (HMDP), previously studied for a variety of cardio-metabolic traits. Metagenomic analysis of cecal DNA followed by genome-wide association analysis identified genomic loci that were associated with microbial enterotypes in the gut. Among these we detected a genetic locus surrounding multiple amylase genes that was associated with abundances of Firmicutes (*Lachnospiraceae* family) and Bacteroidetes (*Muribaculaceae* family) taxa encoding distinct starch and sugar metabolism functions. We also found that lower amylase gene number in the mouse genome was associated with higher gut *Muribaculaceae* levels. Previous work suggests that modulation of host amylase activity impacts the availability of carbohydrates to the host and potentially to gut bacteria. The genetic variants described above were associated with distinct gut microbial communities (enterotypes) with different predicted metabolic capacities for carbohydrate degradation. Mendelian randomization analysis revealed host phenotypes, including liver fibrosis and plasma HDL-cholesterol levels, that were associated with gut microbiome enterotypes. This work reveals novel relationships between host genetic variation, gut microbial enterotypes and host physiology/disease phenotypes in mice.

## Introduction

The microbial communities that inhabit the gut of mammals have profound effects on host biology and health. Alterations in the intestinal microbiome have been associated with myriad conditions including metabolic disorders, cardiovascular disease^1,2^ and immunity^3,4^. Host genetic variation and environmental factors, including diet, modulate the gut microbiome and its interactions with the host^5,6^. Distinct gut microbial communities, termed enterotypes, that reflect stratification in a population and define compositional attributes, have been identified^7,8^. However, it remains largely unknown how host genetics modulates microbial enterotypes as well as enterotype-associated functions and metabolic pathways in the gut.

Carbohydrates represent an important energy source for both human and microbial cells. Dietary compounds that cannot be digested by host enzymes, including plant polysaccharides and resistant starches, reach the distal gut where they are broken down and fermented by resident bacteria^9^. Consumption of different carbohydrates can influence the gut microbiota and its association with the host^10,11^. Non-digestible carbohydrates such as dietary fibers resist host digestion and can therefore serve as substrates for microbiota in the distal gut. Digestible carbohydrates like starch, on the other hand, are broken down by host amylases^12^, releasing its constituent sugars which can be absorbed by the host or used by gut microbes. Amylase gene copy number varies among individual humans and associations between amylase copy number and diet have been reported for mammals^12–14^. In mice, the genes encoding salivary (*Amy1*) and pancreatic (*Amy2b*) amylases are located in a gene cluster on chromosome 3, whereas the copy number of the *Amy2b* paralogous gene (*Amy2a1*, *Amy2a2*, *Amy2a3*, *Amy2a4*, *Amy2a5*) is variable across different mouse strains^13^. Recent work has revealed that humans with higher salivary amylase gene (*AMY1*) copy number harbor gut microbiomes with increased abundance of resistant starch-degrading microbes and produced higher levels of short-chain fatty acids and elicited higher adiposity when transplanted into germ-free animals^15^. This result suggests that genetic variation of host amylase gene locus may potentially impact gut microbiome and its subsequent effects on the host.

Several mouse and human studies have examined the role of host genetics in shaping the composition of the gut microbiota. These efforts mostly used 16S rRNA gene sequencing^16–18^. Shotgun metagenomics allows comprehensive profiling of the functions and metabolic pathways present in gut communities. However, a limited number of studies have applied metagenomic characterization of gut microbiome in large-scale genetically diverse, phenotypically characterized cohorts. A valuable resource for such an undertaking is the Hybrid Mouse Diversity Panel (HMDP), which consists of over 100 common inbred and recombinant inbred strains. In order to study cardio-metabolic traits, each strain from this panel was made hyperlipidemic by transgenic expression of human apolipoprotein E-Leiden (APOE-Leiden) and human cholesteryl ester transfer protein (CETP). This was done by breeding each of the HMDP strains to the C57BL/6J strain that possessed these two hyperlipidemia-inducing transgenes. Genetic differences among these animals arise only from sequence variations present in the individual recipient strains. This set of F1 animals was termed Ath-HMDP and represents a rich resource for studies exploring complex interactions underlying atherosclerosis and liver fibrosis^19–21^. Here, we used shotgun metagenomics to characterize the gut microbiome from 90 Ath-HMDP strains fed a high-fat diet supplemented with 1% cholesterol. Our analysis identified three microbial enterotypes in this mouse cohort, dominated by Firmicutes, Bacteroidetes or Verrucomicrobiota, and identified two host genomic loci that are significantly associated with the enterotypes. Our results suggest that genetic variation, potentially in host amylase genes, could be a determinant of gut microbial enterotypes by selecting specific carbohydrate metabolizing bacterial species. Furthermore, our results suggest that the functional niches of the selected enterotypes subsequently influence host biomarkers including plasma cholesterol and triglycerides, and disease phenotypes including liver fibrosis.

## Results

### Characterization of the gut metagenomes from 90 Ath-HMDP mouse strains

We generated metagenomic datasets from DNA extracted from cecal contents collected from 356 F1 Ath-HMDP mice encompassing 90 strains (males and females were included for most strains) fed a high-fat diet (33% kcal from fat) with 1% cholesterol for 16 weeks (37.8 million paired-end reads/sample). The generation of this mouse cohort has been previously described^19^. Metabolic and disease-associated phenotypes including plasma lipids, glucose levels, atherosclerotic lesions development, and liver fibrosis varied widely among strains^19,20^ (**Supplementary Table 1**).

Phylogenetic and functional analyses of these metagenomes identified 461 bacterial taxa (7 phyla, 43 classes, 49 orders, 63 families, 131 genera and 166 species), 2,127 KO (KEGG Orthology) functions and 294 metabolic pathways (MetaCyc database) across all mice. Gut microbiota composition was highly variable across the 90 strains; for example, the relative abundance of Firmicutes phylum ranged from 10% to 55% whereas the relative abundance of Bacteroidetes ranged from 5% to 77% (**Supplementary Fig. 1A**). The most abundant species (> 0.5%) that were present in at least 90% of mice included the *Paramuribaculum intestinale*, *Muribaculum intestinale*, *Muribaculum gordoncarteri, Duncaniella muris, Duncaniella freteri*, *Alistipes finegoldii*, *Parabacteroides goldsteinii*, *Faecalibaculum rodentium*, *Lachnospiraceae bacterium*, *Bilophila sp.* and *Parasutterella sp* (**Supplementary Fig. 1B, Supplementary Table 2**). For the purpose of our analyses, we defined these taxa as the “core microbiome species” of the Ath-HMDP mice.

### Gut enterotypes of the Ath-HMDP microbiome

Principal coordinate analysis (PCoA) of the microbial communities described above using Bray-Curtis distance of species abundance resulted in three clusters each dominated by a different phylum: (i) Firmicutes, (ii) Bacteroidetes and (iii) Verrucomicrobiota respectively (**Supplementary Fig. 2A**). Using microbial function abundance resulted in similar clustering each dominated by one of these three phyla (**Supplementary Fig. 2B**). The first principal component (PC1) of the microbial functions, which explains the most variance for the functional profiles, was highly correlated with several taxa, including the *Lachnospiraceae* family (Spearman’s 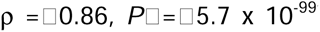) and the *Desulfovibrionaceae* family (Spearman’s ρ 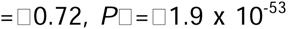) (**Supplementary Table 3**). Because we observed a stratification in the taxonomic composition and functions of the gut microbiome, we next explored the existence of different enterotypes. Using partitioning around medoid (PAM) clustering of Bray-Curtis distance of species abundance, we detected three enterotypes, each of which was identifiable by the levels of Firmicutes, Bacteroidetes and Verrucomicrobiota respectively (**Fig. 1A-B**). Previous studies reported distinct enterotypes dominated by *Bacteroides*, *Prevotella* and *Ruminococcaceae* in the human gut^7,8^. We observed clusters dominated by different taxa as compared to humans, which is likely due to the distinct overall gut microbiota composition detected in these two mammals.

**Fig. 1.**
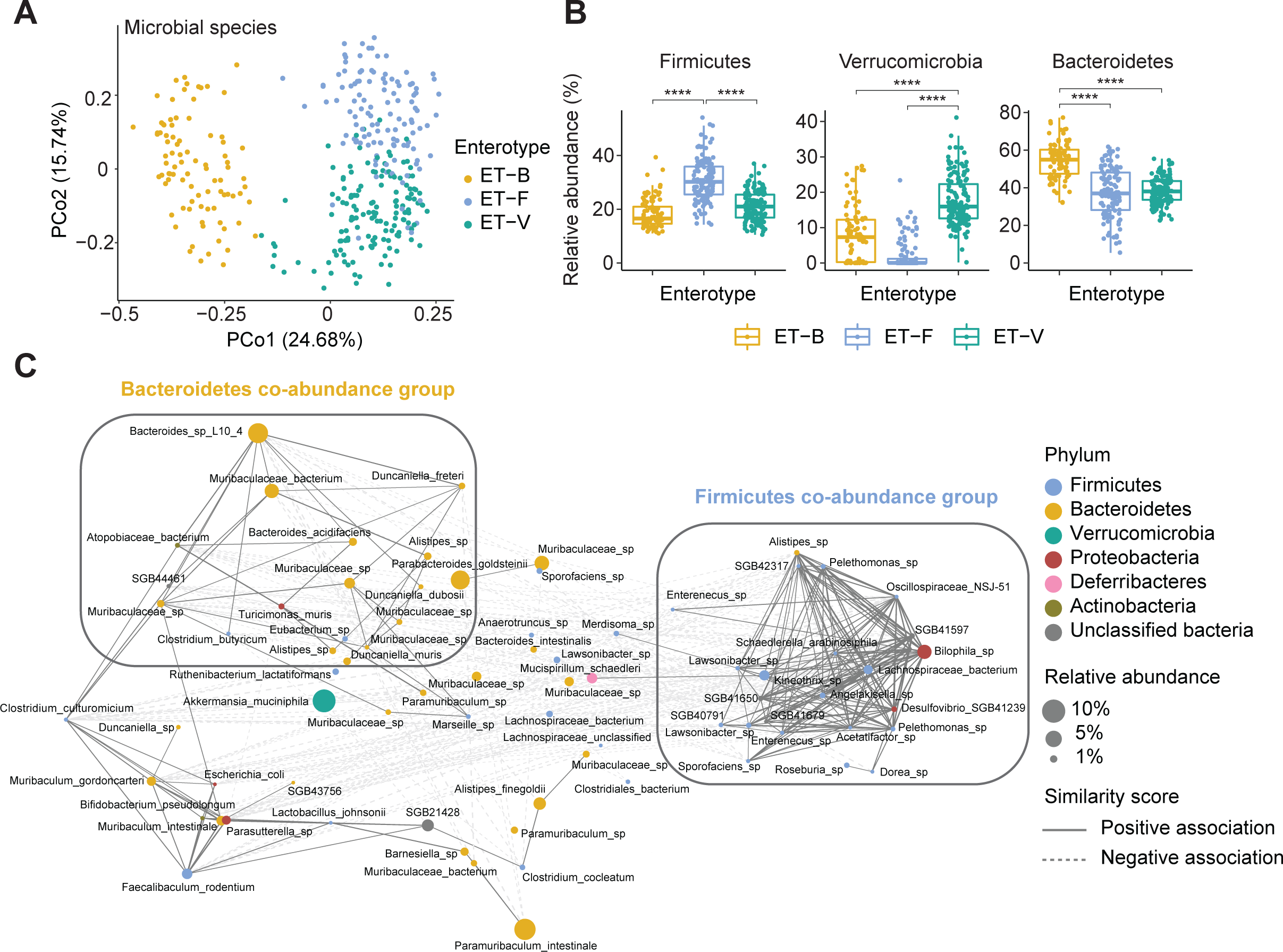
Enterotypes and bacterial species co-abundance groups. **A.** Clustering identified three gut microbial enterotypes in the Ath-HMDP mouse cohort. **B.** Relative abundance of Firmicutes, Bacteroidetes and Verrucomicrobia in three enterotypes. **C**. Correlation network of bacterial species (average relative abundance > 0.1 % and present in at least 20% of samples) using CCREPE with a checkerboard score, indicating the co-occurrence or co-exclusion between species. Nodes represent the species and lines represent similarity score. Solid lines represent co-occurrence species and dashed lines represent co-exclusion species. Significance was calculated by unpaired two-tailed Welch’s t-test and is designated as follows: ** p value < 0.01; *** p value < 0.001; **** p value < 0.0001. ET-B, Bacteroidetes enterotype; ET-F, Firmicutes enterotype; ET-V, Verrucomicrobia enterotype.

To identify microbial species that shape the enterotypes, we summarized the most abundant and prevalent species (relative abundance > 0.1% and present at least > 20% mice) into co-abundance groups (CAGs) using co-occurrence scores (**Fig. 1C**). The Bacteroidetes-dominated CAG contained mostly Bacteroidetes taxa including *Muribaculaceae*, the most abundant family in mouse gut^22^, (*Duncaniella muris, Duncaniella freteri*, *Duncaniella dubosii*, *Muribaculaceae CAG-495 sp., Muribaculaceae CAG-873 sp.*), *Bacteroides* genus (*Bacteroides acidifaciens*, *Bacteroides sp.*) and *Alistipes sp*. The Firmicutes-dominated CAG consisted mostly of Firmicutes taxa including *Lachnospiraceae* family (*Roseburia sp., Dorea sp., Acetatifactor sp., Sporofaciens sp., Kineothrix sp.* and *Schaedlerella arabinosiphila*), *Oscillospiraceae* family (*Lawsonibacter sp., Enterenecus sp.* and *Pelethomonas sp.*) and Proteobacteria including *Bilophila sp*.

### Gut microbiome features are associated with host genetics

To identify associations between mouse genomic variation and gut microbiome features, we performed genome-wide association analysis and mapped the abundance of bacterial functions, taxa and metabolic pathways to the mouse genome (**Supplementary Fig. 3**). We observed 646 functions, 138 taxa and 109 pathways that were associated with at least one significant locus by genome-wide significance cutoff (*P* < 4 x 10^-6^). We further examined the density of these significantly associated gut microbial traits over the whole mouse genome and identified a GWAS hotspot on chromosome 3 at 113-115 Mbp. This genomic locus was associated with 347 different bacterial functions. Pathway enrichment analysis using Fisher’s exact test revealed that genes encoding bacterial flagellar assembly, bacterial chemotaxis and bacterial motility proteins were significantly overrepresented among the functional traits mapping to this GWAS hotspot (**Fig. 2A**).

**Fig. 2.**
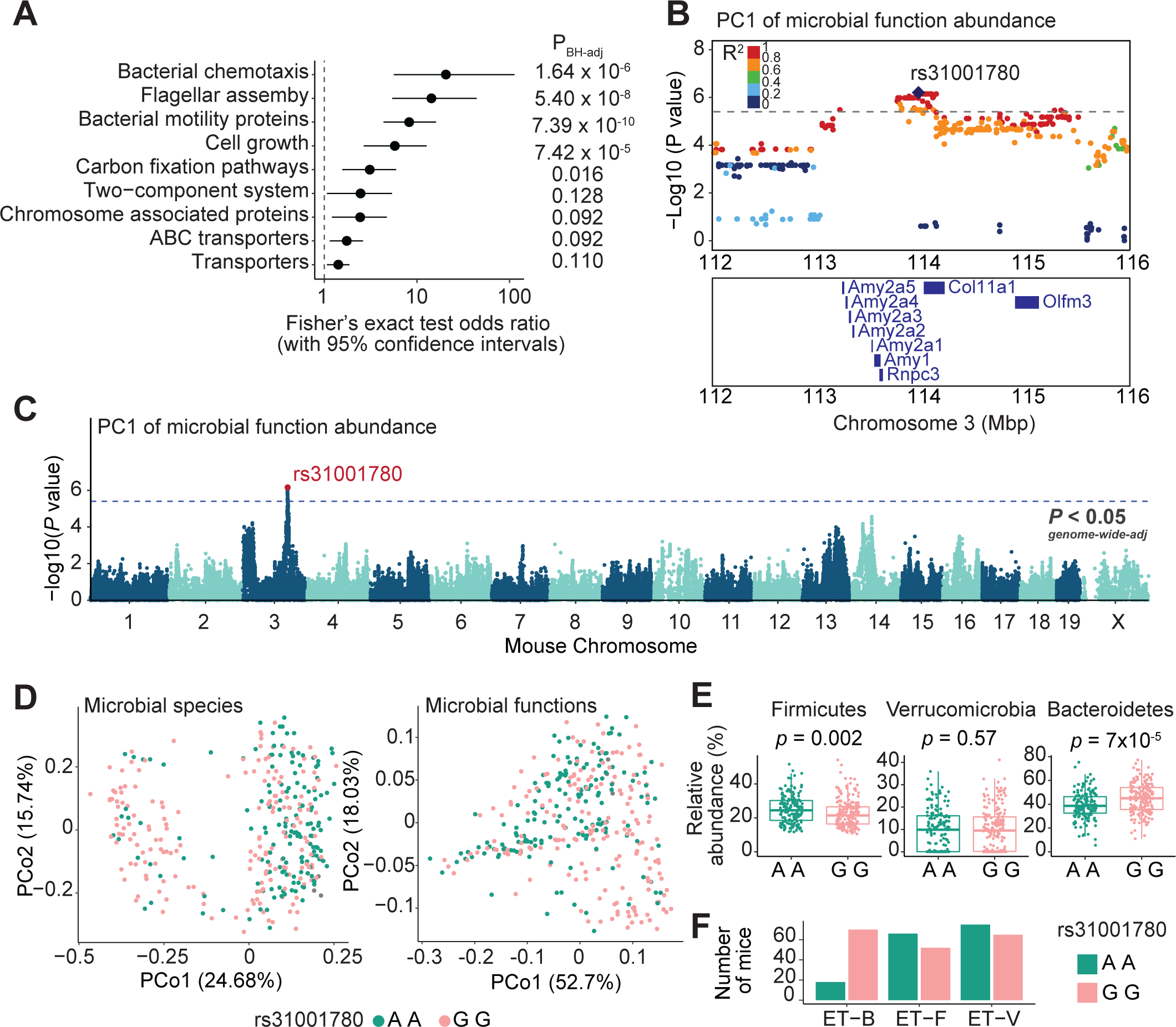
Host genomic loci are associated with gut microbial features. **A.** Enrichment analysis using Fisher’s exact test for gut bacterial functions mapping at the hotspot locus on Chr3: 112– 116lJMbp. **B.** SNP associations for microbial functions PC1 on chromosome 3. Protein coding genes are displayed for Chr3: 112–116lJMbp region. **C**. Genomic locus at Chr3: 113-114 Mbp is associated with PC1 of microbial functions. The lead SNP is an intron SNP rs31001780. Dashed lines represent significance thresholds determined by permutation tests (*P* < 4 x 10^-6^). **D**. Mice with alleles AA, AG or GG at SNP rs31001780 are visualized in PCoA of taxonomy beta-diversity (left) and KO functions beta-diversity (right). **E**. Relative abundance of Firmicutes, Bacteroidetes and Verrucomicrobia from mice with AA or GG at SNP rs31001780. **F**. Number of mice that have AA or GG at SNP rs31001780 in each of three enterotypes.

The genomic regions identified by GWAS most likely contain candidate gene/genes that are in strong linkage disequilibrium (LD) with the lead SNP. Nine protein coding genes are in the LD region (determined by correlation *r*^2^ with lead SNP > 0.8) at chromosome 3 GWAS hotspot, including mouse amylase cluster genes (*Amy1*, *Amy2a1*, *Amy2a2*, *Amy2a3*, *Amy2a4* and *Amy2a5*), *Rnpc3*, *Col11a1* and *Olfm3* (**Fig. 2B**). Gene expression data were available from a subset (n = 4-8 female mice/ strain; 40 strains) of the same Ath-HMDP mice^19^. We conducted correlation analysis between each functions PC1 and genes within the chromosome 3 GWAS hotspot and found significant correlations with *Amy1* and *Rnpc3* gene expression levels (**Supplementary Fig. 4**). In humans, the salivary amylase gene (*AMY1*) copy number is associated with nearby SNPs, structural haplotypes of the amylase locus^23^, and gut microbiome composition^15^. Our data suggest that genomic variants of the amylase gene locus in mice are associated with abundance of particular bacterial functions.

We found that the PC1 of microbial functions abundance mapped to the same hotspot locus, as expected (**Fig. 2C**). PC1 explained the most variance of microbial functions, which are also clustered by the three enterotypes (**Supplementary Fig. 1B**). The lead SNP rs31001780 has two alleles, A and G, which were differ in prevalence between enterotypes (**Fig. 2D**). More specifically, mice with the allele A at SNP rs31001780 had higher Firmicutes and lower Bacteroidetes levels in the gut (**Fig. 2E-F**). The other genetic locus associated with the enterotypes was on chromosome 1 (**Supplementary Fig. 5A**). This locus has the lead SNP rs31965376 with two alleles, A and T, which are associated with *Akkermansia* levels in the gut (**Supplementary Fig. 5B-D**).

### Co-abundance groups (CAGs) are associated with bacterial carbohydrate metabolism

To explore potential bacterial functions and pathways that were modulated by enterotype associated SNPs, we examined the metabolic niche differences between the two identified CAGs. We first compared correlations between functions from starch and sugar metabolism pathway with the species abundance from two CAGs (**Supplementary Fig. 6**). We found that a number of genes that encode glycosidases were associated with Bacteroides species. Glycosidases are enzymes that break down glycosidic linkages to liberate monosaccharides and oligosaccharides of lower molecular weight from carbohydrate substrates. Among those that we found to be significantly correlated with the Bacteroides CAG were amylosucrase (K05341, predicted to degrade sucrose), β−glucosidase (K05350), β−fructofuranosidase (K01193) and oligo−1,6−glucosidase (K01182). Whereas α−amylase (K07405 and K01176, predicted to degrade starch), endoglucanase (K01179, predicted to degrade cellulose), dextranase (K05988, predicted to degrade dextran) and pullulanase (K01200, predicted to degrade pullulan) were associated with Firmicutes species. In addition, bacterial species from the Firmicutes CAG were positively correlated with sugar transport system and phosphotransferase system (PTS) (**Supplementary Fig. 6**).

We reasoned that the variant within the host amylase genes locus could affect amylase activity, which in turn would result in different carbohydrates being available for microbes in the gut. This would lead to distinct gut microbial communities (enterotypes) with different metabolic capacities. To further examine this hypothesis, we characterized the gut microbiome by shotgun metagenomic sequencing from three mouse strains with different amylase gene copies: C57BL/6J (B6) mice have eight amylase genes (*Amy1*, *Amy2a1*, *Amy2a2*, *Amy2a3*, *Amy2a4*, *Amy2a5*, *Amy2b*, *Amy2-ps1*); NZO/HLtJ (NZO) mice have five amylase genes (*Amy1*, *Amy2a1*, *Amy2a2*, *Amy2b*, *Amy2-ps1*); CAST/EiJ (CAST) mice have three amylase genes (*Amy1*, *Amy2a2*, *Amy2-ps1*) (**Fig. 3A**). These mice were fed a high carbohydrate diet (**Supplementary Table 4**) for 12 weeks. We found the abundance of *Muribaculaceae* family was significantly higher in CAST mice compared to those in B6 and NZO (**Fig. 3B**) and the abundance of bacterial α-amylase enzyme (K07405) was higher in CAST compared to those in B6 and NZO (**Fig. 3B**). These results suggested that a lower amylase gene copy number in the mouse genome resulted in increased *Muribaculaceae* abundance in the gut supporting the hypothesis that genetic variation in the amylase gene region is causal for enterotype associated variants on chromosome 3 in mice.

**Fig. 3.**
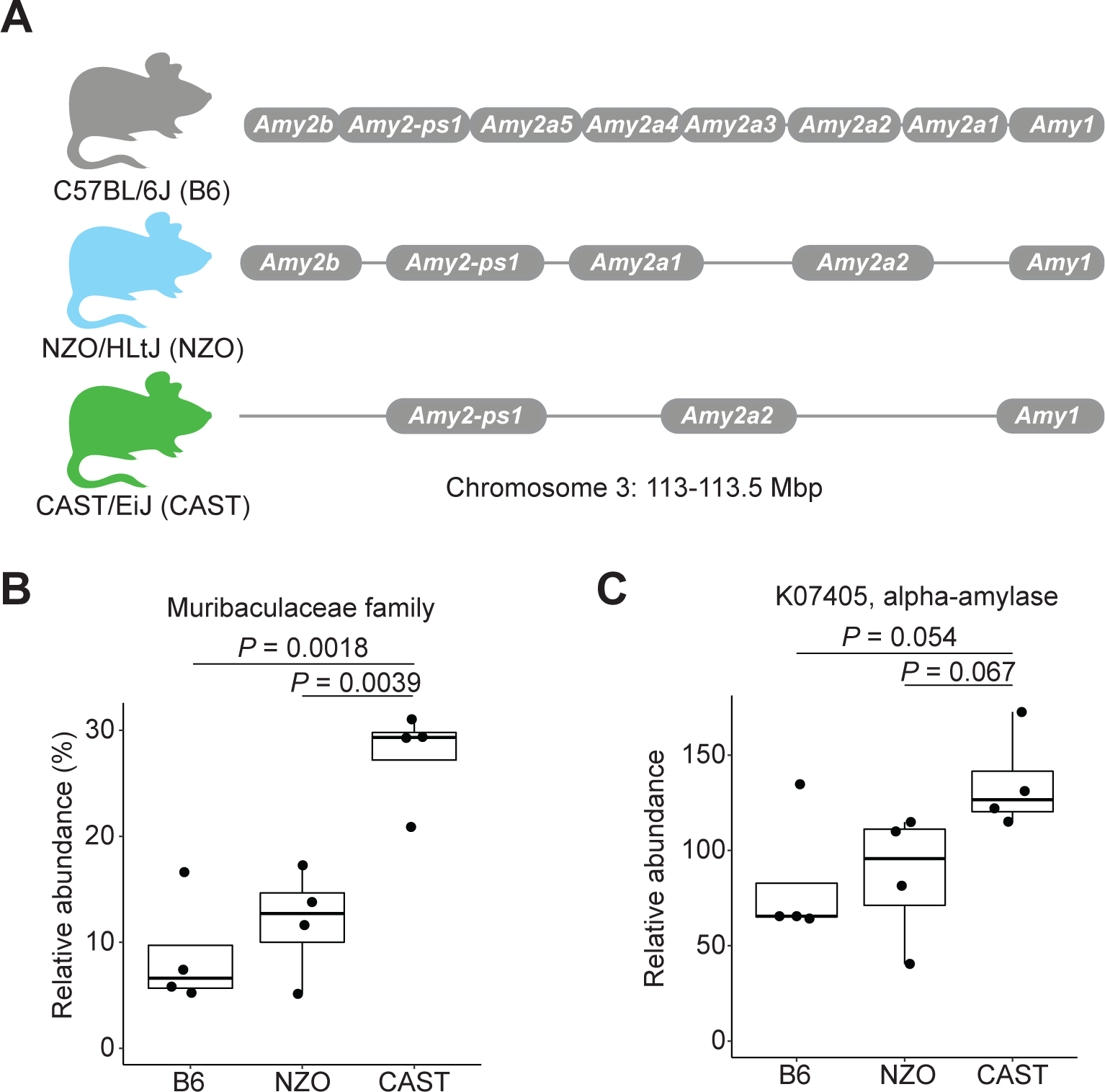
*Muribaculaceae* and gut bacterial alpha amylase genes are more abundant in mice with low amylase copy number. **A.** Amylase gene cluster at chromosome 3 locus from B6, NZO and CAST mouse genome, **B**. Relative abundance of *Muribaculaceae* family in B6, NZO and CAST mice fed on 12 weeks of a high carbohydrate diet. **c**. Relative abundance of bacterial alpha-amylase enzyme (K07405) in B6, NZO and CAST mice fed a high carbohydrate diet for 12 weeks. Statistical difference between treatment groups was tested by unpaired two-sided Welch’s t-test.

### Enterotype species are associated with host traits

We next explored the associations between CAG species and host cardio-metabolic phenotypes for these mice. Species from the two CAGs discussed above showed distinct associations (**Fig. 4A**). Atherosclerotic lesion size was positively correlated with *Bacteroides sp.* (Spearman’s 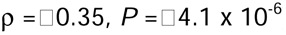) and *Bilophila sp.* (Spearman’s 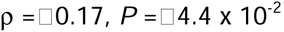) and negatively correlated with *Roseburia sp.* (Spearman’s 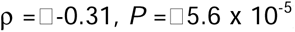), a finding that was also reported in a previous study^26^. We further identified positive associations for *Turicimonas muris* (Spearman’s 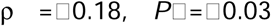), *Atopobiaceae bacterium sp.* (Spearman’s 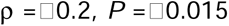) and negative associations for *Dorea sp.* (Spearman’s 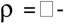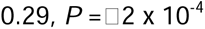) and *Enterenecus sp.* (Spearman’s 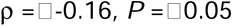) with atherosclerotic lesion size. Plasma levels of HDL and LDL/VLDL were positively correlated with bacteria from Firmicutes CAG and negatively correlated with bacteria from Bacteroidetes CAG. Liver fibrosis was negatively correlated with Bacteroidetes CAG species including *Bacteroides sp.* and taxa within the *Muribaculaceae* family and positively correlated with Firmicutes CAG species such as *Sporofaciens sp*., *Enterenecus sp*. and *Kineothrix sp*. These results suggested that host physiology phenotypes are highly associated with gut microbiome enterotypes (Firmicutes and Bacteroidetes CAG) in mice.

**Fig. 4.**
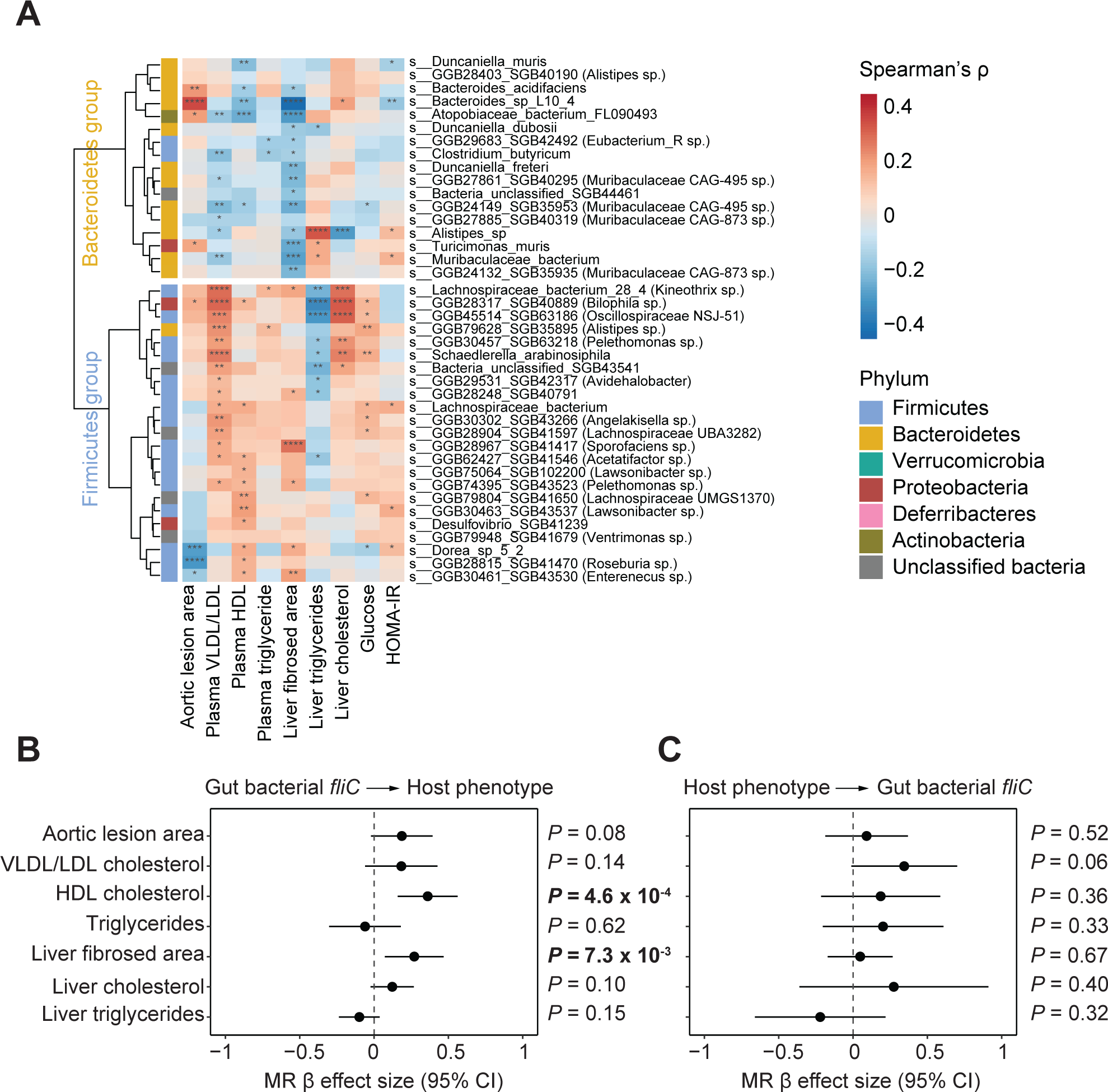
Host phenotypes are associated with CAGs and gut bacteria flagella. **A.** Spearman’s correlation between bacteria species and host physiology phenotypes. Bacteria species from Firmicutes CAG and Bacteroidetes CAG showed different associations. **B**. Bidirectional MR analysis between host clinical traits and gut bacterial flagellin gene abundance. MR results using gut bacterial *fliC* abundance as exposure and clinical traits as outcome. **C**. MR results using clinical traits as exposure and gut bacterial *fliC* abundance as outcome.

### Gut bacterial flagella is associated with host phenotypes

We further examined whether abundance of bacterial flagellar genes was associated with host phenotypes. We focused on bacterial flagellar genes because they were overrepresented among the functions that mapped at chromosome 3 locus and a noteworthy characteristic that distinguish Firmicutes and Bacteroidetes CAGs (**Supplementary Fig. 7**). We observed a strong association between *fliC* abundance (K02406; encoding flagellar filament structural protein) and LDL+VLDL cholesterol (Spearman’s ρ= 0.31, *P* = 5.8 x 10^-10^), HDL cholesterol (Spearman’s ρ = 0.17, *P* = 1.8 x 10^-3^), liver fibrosis area (Spearman’s ρ = 0.17, *P* = 2.5 x 10^-3^), liver cholesterol levels (Spearman’s ρ = 0.24, *P* = 4.1 x 10^-5^) and liver triglycerides (Spearman’s ρ = −0.27, *P* = 4.3 x 10^-6^) (**Supplementary Fig. 8A**). We also detected that *fliC* abundance was correlated with aortic lesion area size in male mice (Spearman’s ρ = 0.2, *P* = 0.052). Sequence polymorphisms in flagellin gene can result in structural variants that influence host recognition and immune responses^27,28^. We examined the most abundant (relative abundance > 0.01%) flagellin gene variants from the Lachnospiraceae (*Roseburia*, *Dorea*) and Desulfovibrionaceae (*Desulfovibrio*) families and found that variants in the nD1 TLR5 epitope motif in the *fliC* gene were associated differently with host physiology phenotypes (**Supplementary Fig. 8B**).

We next performed bi-directional Mendelian randomization (MR) to assess whether gut bacterial flagella causally contribute to host traits. We focused on gut bacterial flagellin (*fliC*) abundance and observed significant causal relationships between gut bacterial *fliC* and liver fibrosed area (*P* = 7.3 x 10^-3^) and HDL cholesterol (*P* = 4.6 x 10^-4^) (**Fig. 4B**). When we tested MR considering clinical traits as the exposure and *fliC* abundance as the outcome, we did not observe significant causal effects (**Fig. 4C**). These results suggest that flagella may influence the progression of liver disease and, consistent with previous work, show that genes involved in flagellar assembly were enriched in fecal microbiomes of patients with moderate to severe fibrosis^29^.

## Discussion

Genome-wide association studies have identified multiple host genomic loci associated with the gut microbiome in humans and mice; however, most of those efforts have focused on organismal composition and there is limited evidence linking specific functions and pathways with host genetic variation^16^. Additionally, gut microbiome enterotypes were described in human cohorts where a small number of bacterial taxa determines the stratification of whole community, but there was limited evidence of enterotypes in genetically diverse mouse cohorts^30^. Our work comprehensively characterized gut microbiome composition, functions, and metabolic pathways in the Ath-HMDP mouse cohort. We identified three enterotypes dominated by different phyla including Firmicutes, Bacteroidetes and Verrucomicrobiota. We also found that the enterotypes were associated with bacterial taxa (Firmicutes and Bacteroidetes CAGs) and microbial functions (starch and sugar metabolism and flagellar assembly). The Bacteroidetes CAG included many bacteria from *Muribaculaceae*, the most abundant family in mouse gut^22^.

Using genetic mapping, we identified host genomic loci associated with bacterial taxa, functions and pathways, and two of these loci were associated with enterotypes. The genetic variant rs31001780 (A/G) at Chr3 locus was significantly associated with Firmicutes and Bacteroidetes enterotypes and genetic variant rs31965376 (A/T) at Chr1 locus was significantly associated with the Verrucomicrobiota enterotype. Importantly, expression level of the *Amy1* gene, which spans in LD region of Chr3 locus, was correlated with Firmicutes, Bacteroidetes and Verrucomicrobiota abundances. Given the same carbohydrate-rich diets, mouse genetic variants of *Amy1* gene may induce different carbohydrate accessibility for gut microbiota and shape the different enterotypes. In humans, individuals with higher salivary amylase gene (*AMY1*) copies show higher levels of many genera within the Firmicutes in the gut^15^, which aligns with our results in mice (i.e., *Amy1* gene expression is positively associated with Firmicutes abundance). We reasoned that the differences in sugar and starch metabolism between Firmicutes and Bacteroidetes CAG species, may explain how genetic amylase locus variants shaped the enterotypes in mouse gut. Previous work showed that mice treated with acarbose, which inhibits amylase activity, have an increased abundance of the *Muribaculaceae* family in gut^24,25^. Members of this family possess starch utilization genes^25^. We found *Muribaculaceae* were more abundant in CAST mice which have a low copy number of pancreatic amylase genes (*Amy2a*). This is the different amylase gene that was identified to be associated with gut microbiome in humans (salivary amylase gene *AMY1*)^15,31^. Both salivary and pancreatic amylases would impact carbohydrate availability and potentially influence the gut microbiome, thus loss-of-function mouse experiments should be conducted to directly test the effects of each amylase on the gut microbiome.

As mentioned above we found genetic variants in the amylase gene locus associated with bacterial flagellin, which was also associated with increased liver fibrosis. Interestingly, previous work showed that treatment with acarbose attenuates experimental non-alcoholic fatty liver disease^32,33^ and non-alcoholic steatohepatitis in mice^34^. Additionally, flagellated Enterobacteriaceae are typically found to be increased in non-alcoholic fatty liver disease (NAFLD) patients with severe fat deposition and fibrosis^35^ and a recent study suggested that *E. coli* promoted the development of NAFLD via flagellin^36^. The causal role of flagella and the mechanism by which it may exacerbate liver fibrosis warrant further investigation.

MR analysis showed that the genes for bacterial flagellin were associated with increased liver fibrosed area and HDL cholesterol levels. Previous studies showed that gut microbiome partially explained variations of plasma triglyceride and HDL cholesterol levels in humans^37^. Another study showed that high-fat diet increased flagellated bacteria in the gut, which increased apolipoprotein A1 (ApoA1) production and HDL cholesterol levels in mice^38^. MR seeks to infer causal effects of modifiable exposure using measured variation in genes of known function. Successful applications of MR in humans have revealed relationships between gut microbiome and other molecular traits, including blood metabolites^39^, short-chain fatty acids^40^, and host metabolic traits^41,42^. To the best of our knowledge, our study is the first MR application of gut microbiome in genetically diverse mouse cohort. Furthermore, our MR results provides evidence for the casual relationship between gut flagellated bacteria and plasma HDL cholesterol levels.

We further reasoned that not only high-fat diet could increase flagellated bacteria in the gut, but the amylase gene copy number can also perhaps indirectly affect the abundance of flagellated bacteria. A recent study showed that bacteria flagellin gene variants from Lachnospiraceae family were associated with differential TLR5 activation^27^. We also found that variants in the nD1 TLR5 epitope motif in the *fliC* gene were associated differently with host physiology phenotypes, including atherosclerotic lesion. This underscores the importance of bacterial genetic variation in gut microbiome association studies. Recent studies have linked bacterial SNPs in human gut microbiome with host phenotypes^43^, but investigations of bacterial SNPs in the mouse gut microbiome have been limited by a lack of host genetic diversity. We demonstrate here that using a genetically diverse cohort of mice, such as HMDP, is a feasible approach to achieve a better understanding of gene-level microbiome associations with the host.

Together, our work highlights how host genetics can shape different microbial enterotypes in the mouse gut and identifies potential candidate host genes involved. The presented results also suggest that the microbial enterotype associated functions and pathways that are the consequence of host genetic variants influence host cardio-metabolic phenotypes.

## Methods

### HMDP mouse cohort

Male and female mice from the F1 Ath-HMDP panel were maintained in a temperature and humidity controlled environment under a 12 h light/dark cycle (lights on at 6:00 and off at 18:00). Mice were housed by strain. All mice were fed a high fat diet (33 kcal % fat from cocoa butter) supplemented with 1% cholesterol (Research Diets D10042101) for 16 weeks. Cecal contents were collected immediately after animals were euthanized. Clinical phenotypes for this cohort were described in a previous study^19^.

### Validation mouse cohort

Male C57BL/6J, NZO/HLtJ and CAST/EiJ mice were maintained in a temperature and humidity controlled environment under a 12 h light/dark cycle (lights on at 6:00 and off at 18:00). Mice were single housed. All mice were fed a high carbohydrate and low fat diet (Envigo Teklad TD.200339) for 12 weeks. Cecal contents were collected immediately after animals were euthanized.

### Metagenomic sequencing

Cecal DNA was extracted from individual mice using the PowerSoil DNA Isolation Kit. DNA concentration was verified using the Qubit® dsDNA HS Assay Kit (Life Technologies, Grand Island, NY). Samples were prepared using Illumina NexteraXT library preparation kit. Quality and quantity of the finished libraries were assessed using an Agilent bioanalyzer and Qubit® dsDNA HS Assay Kit, respectively. Libraries were standardized to 2nM. Paired end, 150 bp sequencing was performed using the Illumina NovaSeq6000. Images were analyzed using the standard Illumina Pipeline, version 1.8.2.

### Profiling microbiome composition

Gut microbial taxon was profiled by MetaPhlAn4 pipeline (ver 4.0.2) using the MetaPhlAn database (mpa_vOct22) and the ChocoPhlAn pan-genome database (mpa_vOct22_CHOCOPhlAnSGB_202212) that contains a collection of around 1lJmillion prokaryotic metagenome-assembled genomes^44^. The taxonomy clades with average relative abundance > 0.01% and present in at least >20% samples were kept as microbial taxon for downstream analyses. The unclassified SBG taxa were further annotated to Genome Taxonomy Database (GTDB) using mpa_vOct22_CHOCOPhlAnSGB_202212_SGB2GTDB.tsv data from MetaPhlan4 pipeline.

### Profiling microbiome function

Raw reads quality control was performed using Trimmomatic (ver. 0.39) with default parameters. To identify and eliminate host sequences, reads were aligned against the mouse genome (mm10/GRCm38) using Bowtie2 (ver. 2.3.4) with default settings and microbial DNA reads that did not align to the mouse genome were identified using samtools (ver. 1.3; samtools view -b -f 4 -f 8). Samples with total read depth <10 million were excluded for downstream analyses. Quantification of microbial genes was done by aligning clean paired end reads to a previous published mouse gut microbiome non-redundant gene catalog^45^ using Bowtie2 (ver. 2.3.4) and default parameters. RSEM (ver. 1.3.1) was used to estimate microbial gene abundance. Relative abundance of microbial gene counts per million (CPM) were calculated using microbial gene expected counts divided by gene effective length then normalized by the total sum. To obtain abundance information for microbial functions, CPM of genes with the same KEGG orthologous (KO) annotation were summed together. In case there were multiple KO annotations for a single gene, we used all KO annotations.

### Profiling microbiome pathway

Gut microbial pathways were profiled using the HUMAnN3 pipeline (ver 3.0.0), the MetaPhlAn database (mpa_v20_m200), the ChocoPhlAn pan-genome database (v296_v201901b) and the UniRef90 protein database (ver 0.1.1)^46^. Pathways with average relative abundance > 0.01% detected in at least >20% samples were used for downstream analyses.

### Enterotype clustering

Gut microbiome data was clustered using partitioning around medoid (PAM) clustering via pam() function from R package cluster (ver. 2.1.2). The Bray-Curtis distance of species abundance was used for PAM clustering. The dominant phylum in each cluster was determined by highest abundant taxon comparing to other clusters.

### Microbial co-abundance groups

Similarity scores between species were calculated using CCREPE (compositionality corrected by renormalization and permutation) package (ver. 1.1.3)^47^. The species network was visualized using ggnet2() function from R package ggnet (ver. 0.1.0).

### Genome-wide association of gut microbiome

The Mouse Diversity Genotyping Array was used for genotyping and gave approximately 450,000 SNPs. SNPs used for each trait were filtered by the following criteria: the minor allele frequency (MAF) > 5% and missing genotype frequency < 10%. GWAS analysis was performed using FaST-LMM^48^ (Python ver. 3.7.4). When testing SNPs on chromosome N, all SNPs from other chromosome besides N were used for kinship matrix construction, that is leave out one chromosome (LOOC) approach. Sex was used as covariate in the regression model. All individual mice were included in GWAS regression model. GWAS significance thresholds were determined by permutation tests^19^. A genome-wide significance threshold of *P* < 4 x 10^-6^ was used. We defined a study-wide significance threshold of *P* < 4 x 10^-6^/ (2127+108+300) = 1.58 x 10^-9^. Broad sense heritability for each trait was estimated using “repeatability()” function from “heritability” (ver. 1.3) R package. Narrow sense heritability for each trait was calculated using all filtered SNPs to estimate the proportion to explain total variations for each trait.

### MR analysis

Bidirectional MR analysis were performed to first test if microbiome traits causally affect a host phenotype and then test if the host phenotypes can causally affect the microbiome traits. We identified independent genetic variants with *P* < 1 x 10^-4^ as instrument variables in MR. We carried out inverse variance weighted (IVW) test using R package TwoSampleMR^49^ (ver. 0.5.6).

### Data and statistical analysis

All data integration and statistical analysis were performed in R (v3.6.3). Differences between groups were evaluated using unpaired two-tailed Welch’s t-test. Enrichment analysis was performed with Fisher’s exact test using a custom R function. Correlation analysis was performed with two-sided Spearman’s correlation using the R function ‘cor.test()’. For multiple testing, Benjamini-Hochberg false discovery rate (FDR) procedure was used to adjust P values. Data integration was performed using R packages dplyr (v1.0.6), tidyr (v1.1.3), reshape2 (v1.4.4) and data.table (v1.14.0). Heat maps were plotted using the R package pheatmap (v1.0.12). Other plots were created using the R packages ggplot2 (v3.3.3).

## Data availability

Shotgun metagenomic data are available from the Sequence Read Archive (SRA) under accession PRJNA1078419.

## Code availability

All code used in this study is available in GitHub (https://github.com/qijunz/Zhang_HMDP_paper) or in the corresponding software package websites.

## Acknowledgements

We thank the University of Wisconsin Biotechnology Center DNA Sequencing Facility for providing sequencing and support services; the University of Wisconsin Center for High Throughput Computing (CHTC) in the Department of Computer Sciences for providing computational resources, support and assistance; This work was supported by National Institutes of Health (NIH) grants HL144651 (F.E.R., A.J. L), HL148577 (F.E.R., A.J. L) and RC2DK125961 (A.D.A.). E.R.H. and M.F.W. were supported in part by the Metabolism and Nutrition Training Program NIH T32 (DK007665). E.R.H. was supported by the University of Wisconsin–Madison Food Research Institute (Robert H. and Carol L. Deibel Distinguished Graduate Fellowship in Probiotic Research). This work was also supported by a grant from a Transatlantic Networks of Excellence Award from the Leducq Foundation (17CVD01).

## Contributions

F.E.R. and A.J.L. conceived the study. Q.Z. designed the experiments. Q.Z. and E.R.H. contributed to sample processing for DNA sequencing. Q.Z. performed metagenomic, network and GWAS analyses. C.P. assisted with GWAS analyses. M.F.W., M.P.K. and A.D.A. assisted with high carbohydrate mouse experiment. Q.Z. and F.E.R. wrote the manuscript. All authors approved the final manuscript.

## Supplemental information

**Supplementary Table 1. Summary of host physiologic traits and gut metagenome in Ath-HMDP mice.** Host cardio-metabolic phenotypes in Ath-HMDP mice were obtained from previous studies. Gut metagenomes were characterized from 356 Ath-HMDP mice, encompassing 90 strains (190 male mice and 166 female mice).

**Supplementary Table 2. Gut microbiome composition traits in the Ath-HMDP mice using MetaPhlAn4.**

**Supplementary Table 3. Spearman’s correlation between the first Principal Component (PC1) of KO profile with bacteria taxa in gut.**

**Supplementary Table 4. High carbohydrate diet use in validation studies.**

---

**Supplementary Fig. 1.**
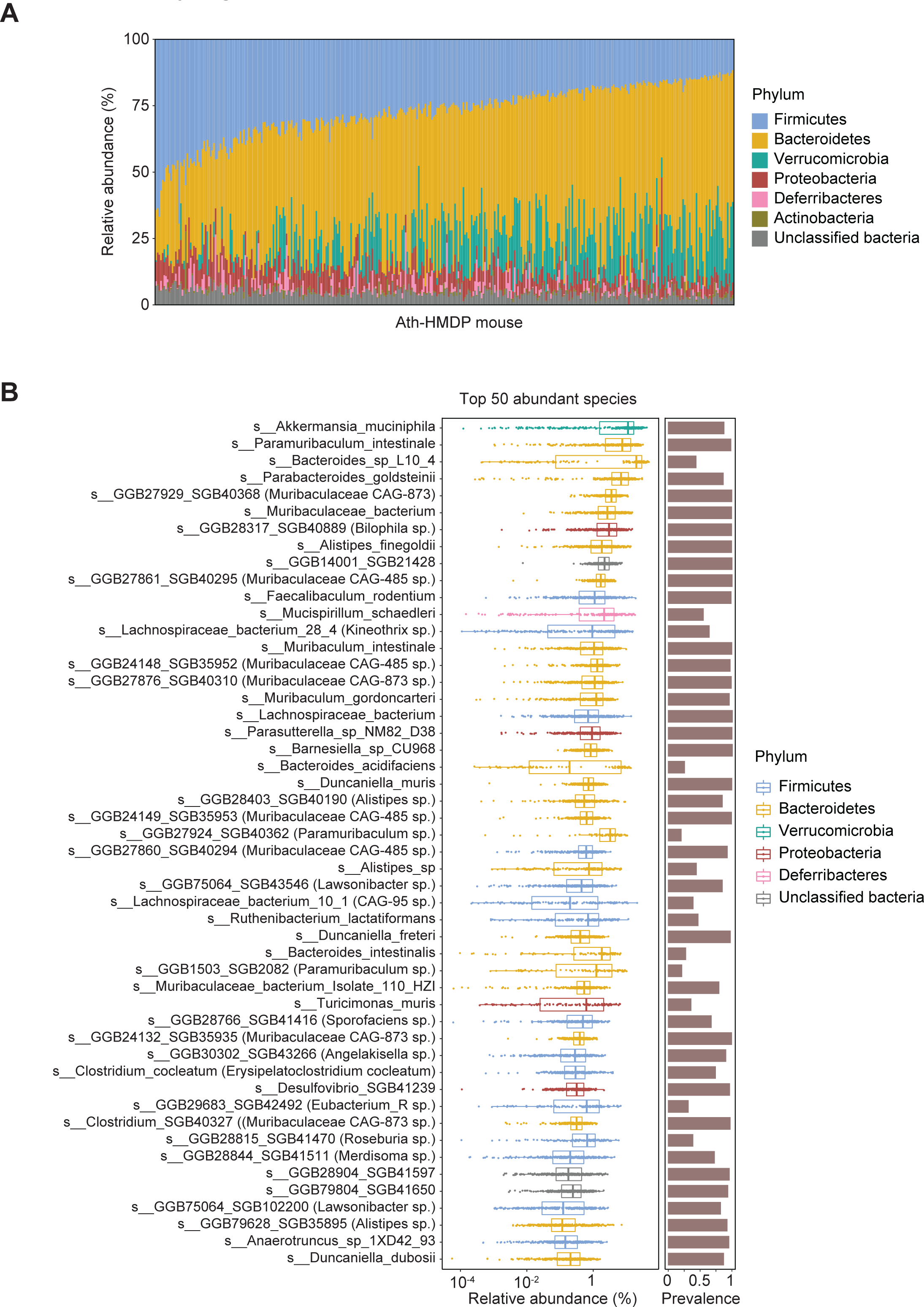
Gut microbiota composition summary. **A**. The relative abundance of phyla level taxa in Ath-HMDP mouse. **B**. The relative abundance of top 50 bacteria species and their prevalence in Ath-HMDP mouse cohort.

**Supplementary Fig. 2.**
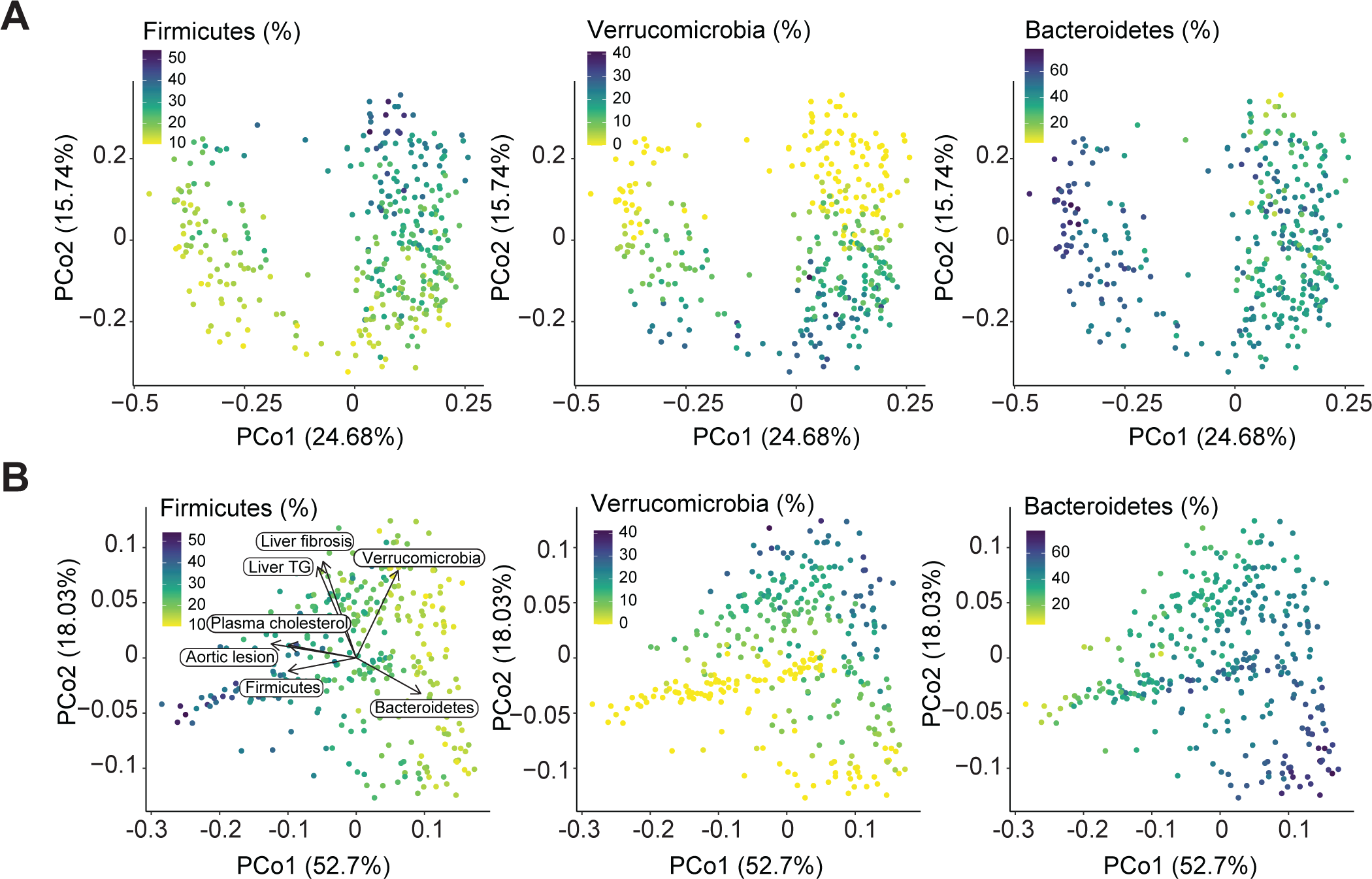
Gut microbial species and functions in the Ath-HMDP mice. **A**. PCoA showing community-level difference among mice in the cohort. Three major phyla show large inter-individual variance and stratify the mouse samples. **B.** PCoA visualizing the beta-diversity of functions in the cohort. Three major phyla show large inter-individual variance and stratify the mouse samples.

**Supplementary Fig. 3.**
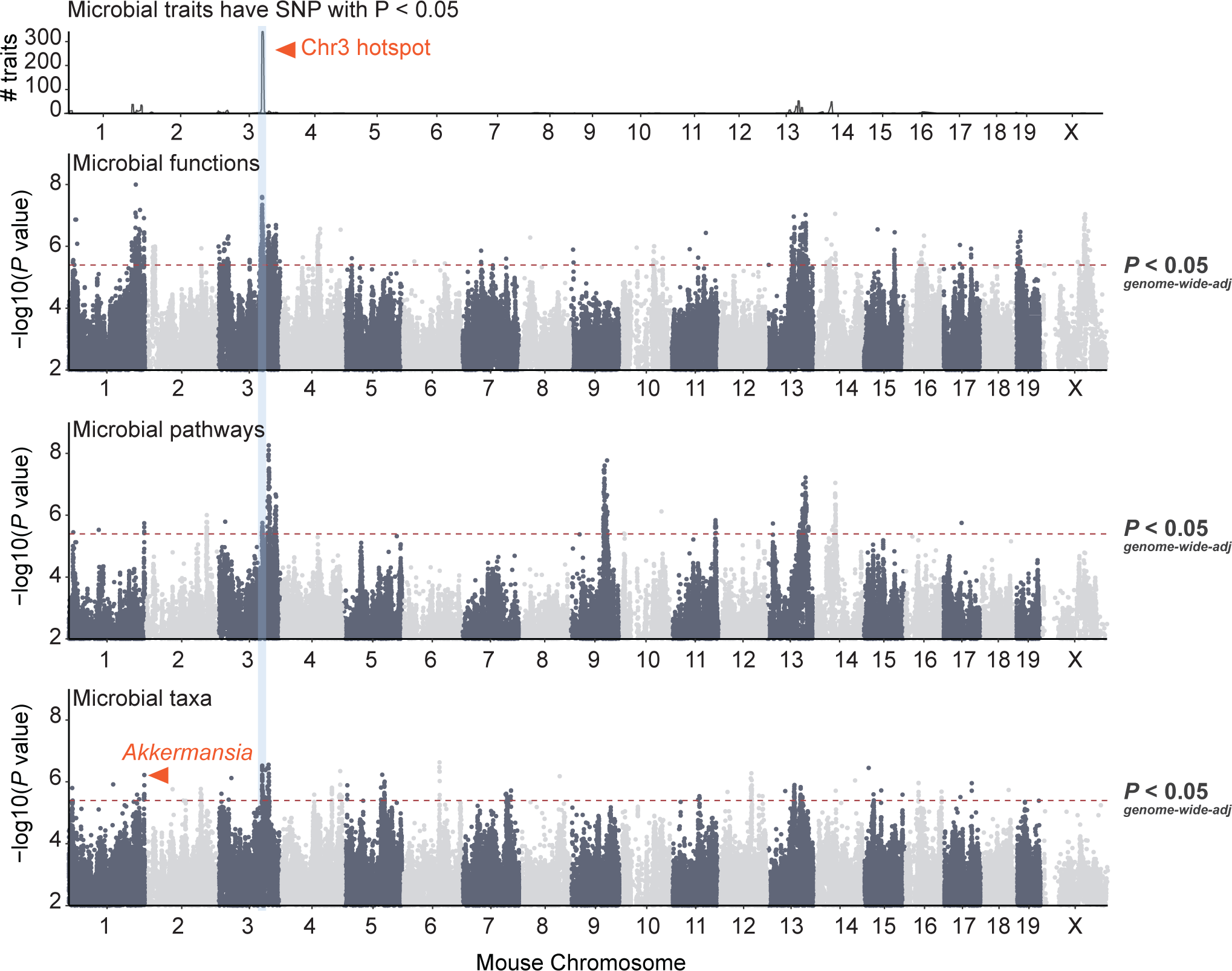
Genetic associations of gut microbial features in Ath-HMDP mice. Genetic associations of gut microbial functions, taxa and metabolic pathways. The density of associated gut bacterial functions in the mouse genome is showed on the top. Dashed lines represent significance thresholds determined by permutation tests (*P* < 4 x 10^-6^).

**Supplementary Fig. 4.**
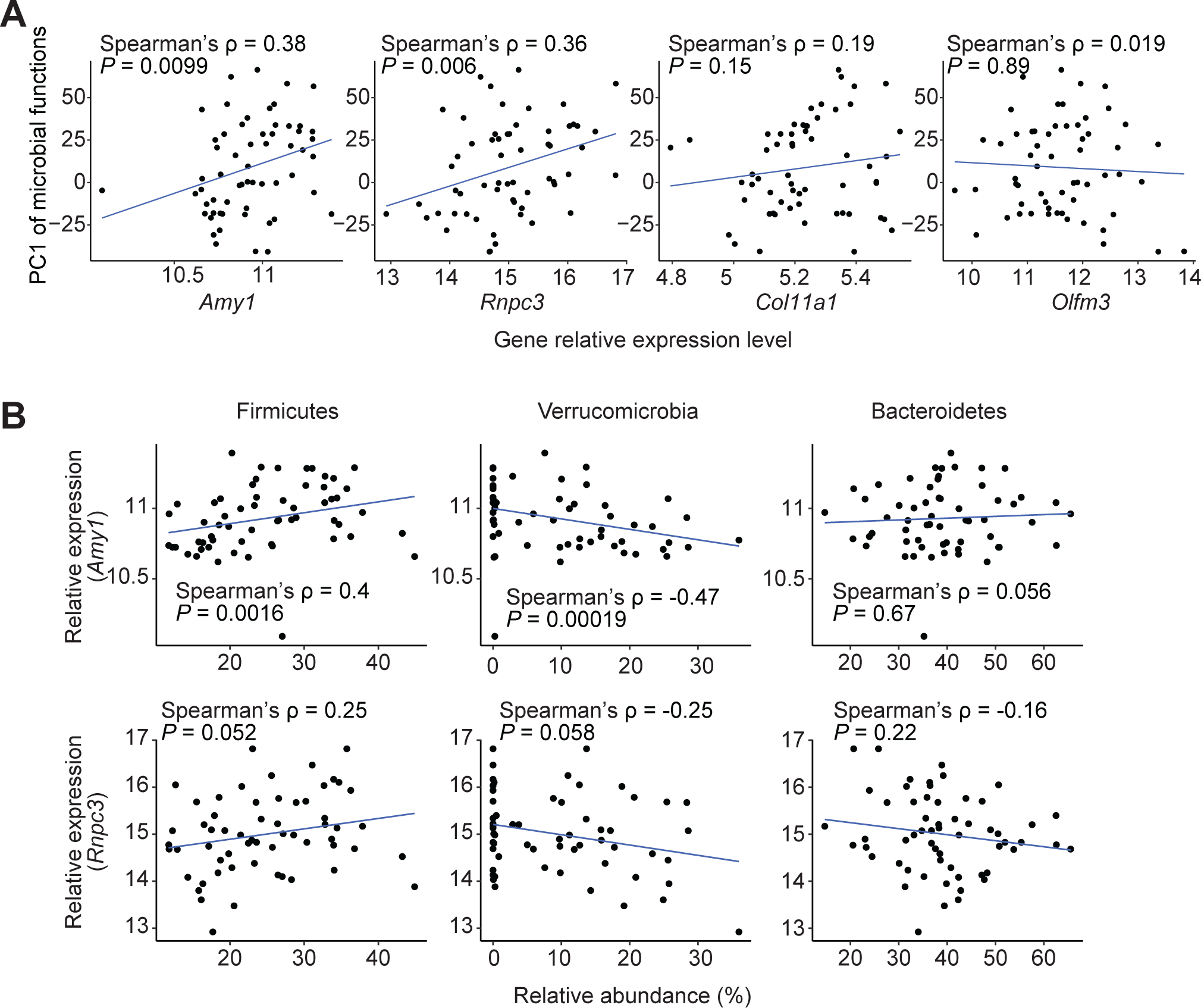
*Amy1* gene expression level correlates with microbial enterotypes. **A**. Spearman’s correlation between candidate genes (*Amy1*, *Rnpc3*, *Col11a1* and *Olfm3*) in Chr3 locus with PC1 of microbial functions. **B**. Spearman’s correlation between candidate genes (*Amy1* and *Rnpc3*) in Chr3 locus with abundance of Firmicutes, Verrucomicrobia and Bacteroidetes.

**Supplementary Fig. 5.**
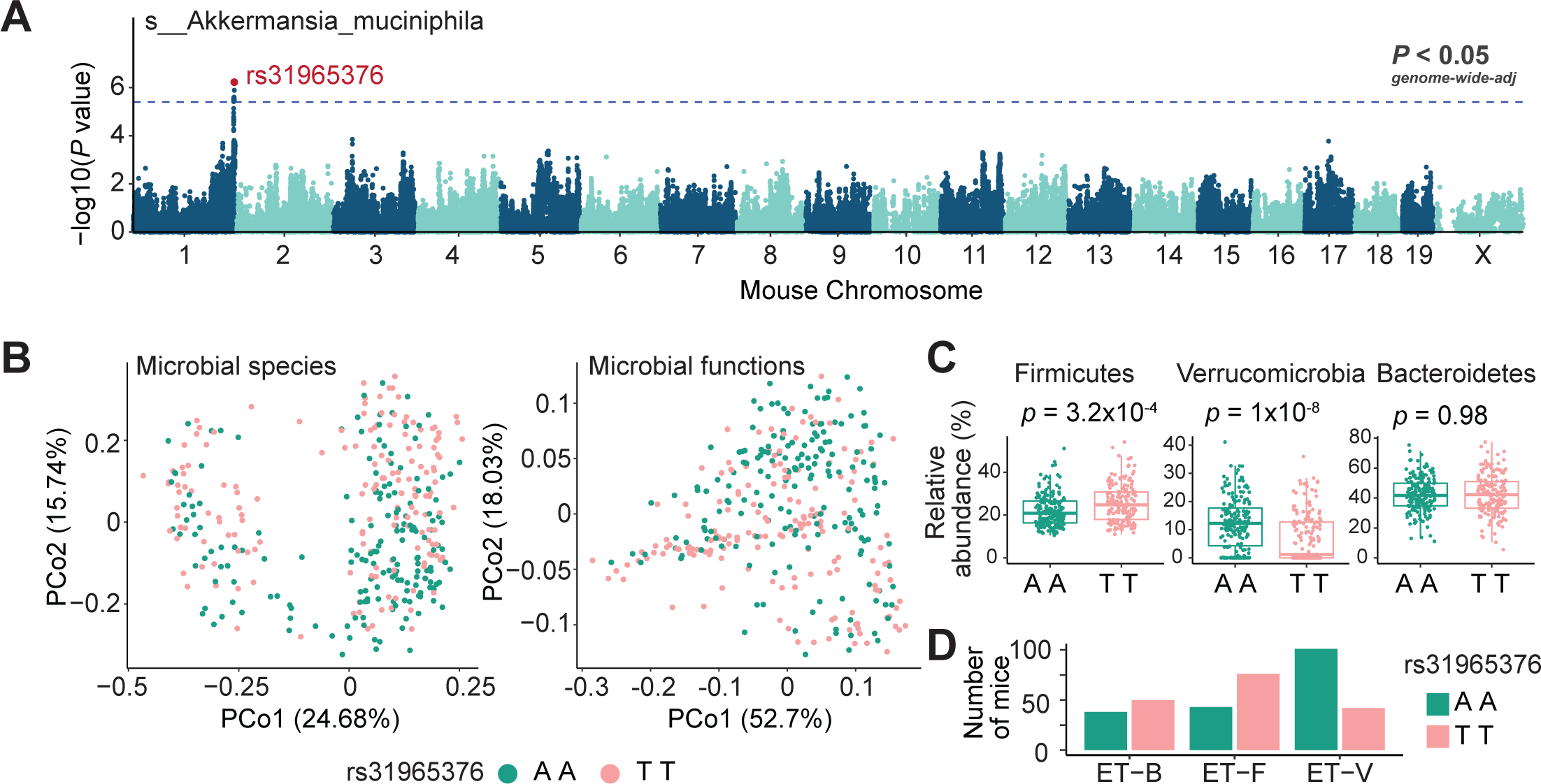
Genomic loci are associated with gut *A. muciniphila* abundance. **A**. Genomic locus at Chr1 193-194 Mbp are associated with *Akkermansia muciniphila* abundance. The lead SNP is rs31965376. Dashed lines represent significance thresholds determined by permutation tests (*P* < 4 x 10^-6^). **B**. Mouse with allele AA or TT at SNP rs31965376 visualized in PCoA of species beta-diversity (left) and functions beta-diversity (right). **C**. Relative abundance of Firmicutes, Bacteroidetes and Verrucomicrobia from mouse with AA or TT at SNP rs31965376. **D**. Number of mice with AA or TT at SNP rs31965376 in each of three enterotypes.

**Supplementary Fig. 6.**
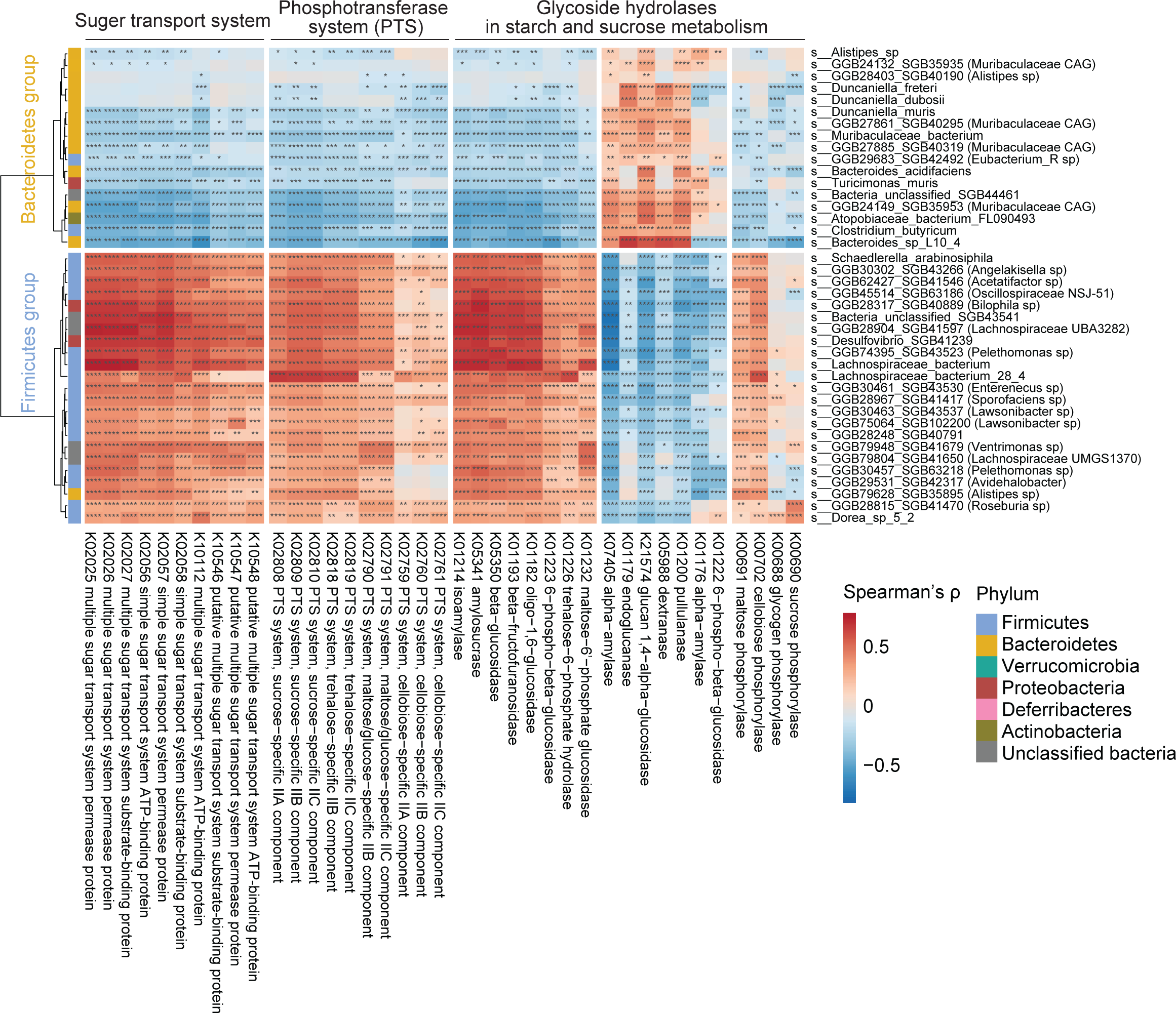
Bacterial starch and sugar metabolism is associated with microbial enterotypes’ species. Spearman’s correlation between bacterial species and bacterial functions involved in starch and sugar metabolism. Bacterial species from Firmicutes CAG and Bacteroidetes CAG showed different associations.

**Supplementary Fig. 7.**
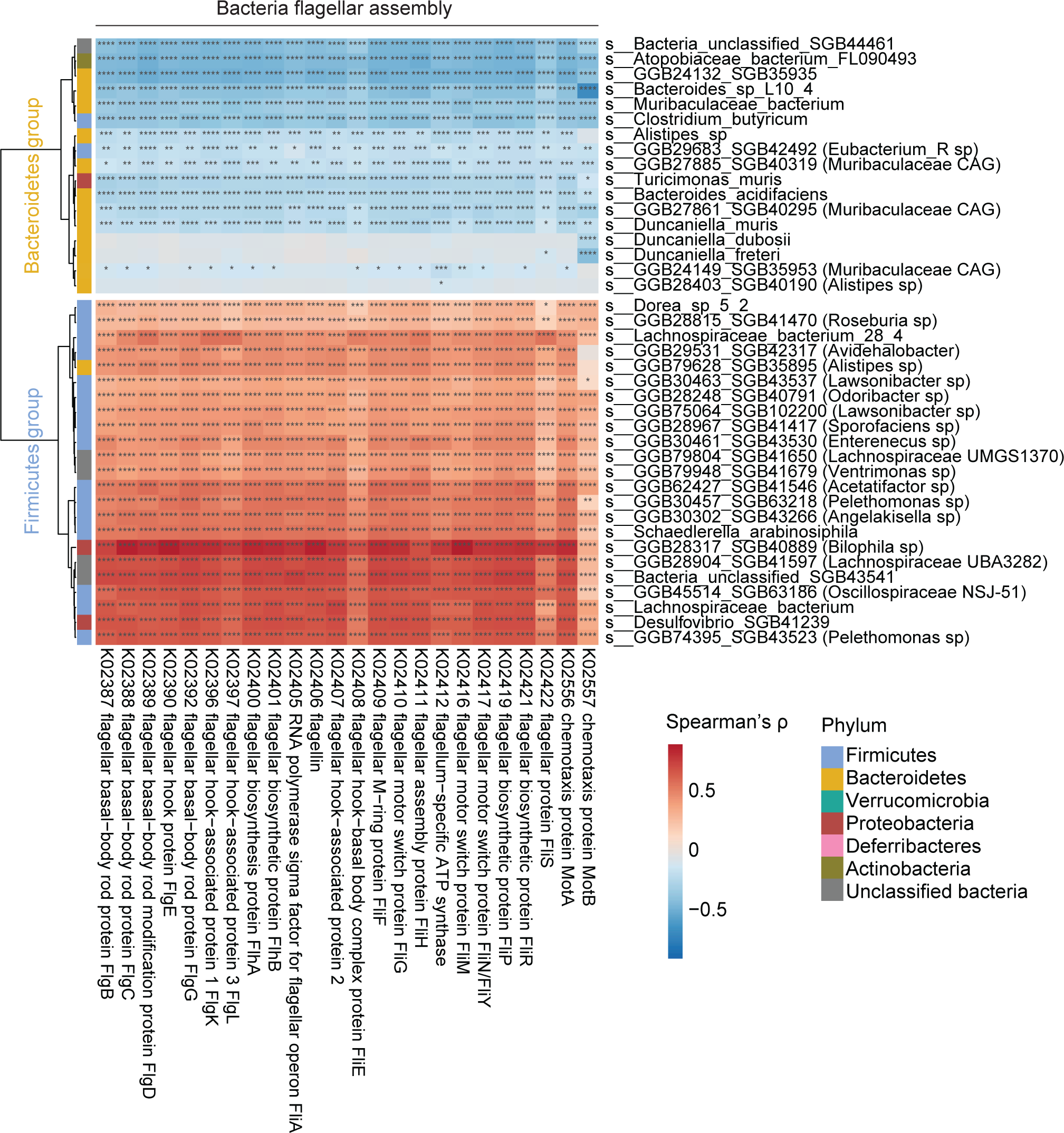
Bacteria flagellar assembly genes are associated with microbial enterotypes species. Spearman’s correlation between bacteria species and bacterial functions involved in flagellar assembly. Bacterial species from Firmicutes CAG and Bacteroidetes CAG showed different associations.

**Supplementary Fig. 8.**
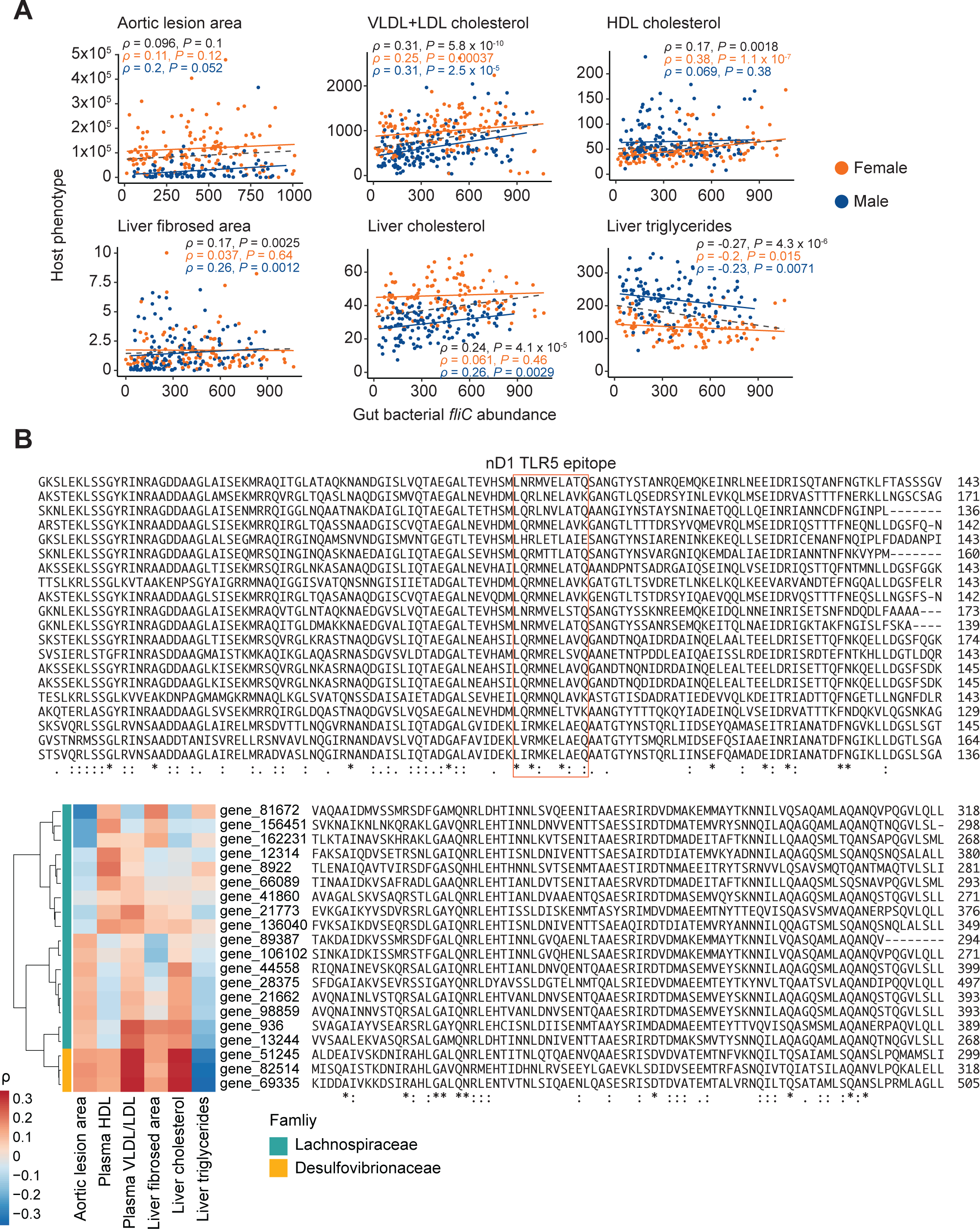
Correlation between gut bacterial fliC gene with host phenotypes. **A**. Spearman’s correlation between host clinical traits and gut bacterial flagellin gene abundance. **B**. Spearman’s correlation between host clinical traits and most abundant individual gut bacterial flagellin gene abundance from Lachnospiraceae and Desulfovibrionaceae family. The flagellin gene sequences were aligned and conserved motifs were identified.

